# Presence of orange tubercles does not always indicate accelerated low water corrosion

**DOI:** 10.1101/855676

**Authors:** Hoang C. Phan, Scott A. Wade, Linda L. Blackall

**Author notes:** Address correspondence to Hoang C. Phan. Present contact: HC Phan, Virbac, Regional RDL Center Asia, Vietnam.

## Abstract

The rapid degradation of marine infrastructure at the low tide level due to accelerated low water corrosion (ALWC) is a problem encountered worldwide. Despite this, there is limited understanding of the microbial communities involved in this process. We obtained samples of the orange-coloured tubercles commonly associated with ALWC from two different types of steel sheet piling, located adjacent to each other but with different levels of localised corrosion, at a seaside harbour. The microbial communities from the outer and inner layers of the orange tubercles, and from adjacent seawater, were studied by pure culture isolation and metabarcoding of the 16S rRNA genes. A collection of 119 bacterial isolates was obtained from one orange tubercle sample, using a range of media with anaerobic and aerobic conditions. The metabarcoding results showed that sulfur and iron oxidisers were more abundant on the outer section of the orange tubercles compared to the inner layers, where Deltaproteobacteria (which includes many sulfate reducers) were more abundant. The microbial communities varied significantly between the inner and outer layers of the orange tubercles and also with the seawater, but overall did not differ significantly between the two steel sheet types. Metallurgical analysis found differences in composition, grain size, ferrite-pearlite ratio and the extent of inclusions present between the two steel types investigated.

**IMPORTANCE:** The presence of orange tubercles on marine steel pilings is often used as an indication that accelerated low water corrosion is taking place. We studied the microbial communities in attached orange tubercles on two closely located sheet pilings that were of different steel types. The attached orange tubercles were visually similar, but the extent of underlying corrosion on the different steel surfaces were substantially different. No clear difference was found between the microbial communities present on the two different types of sheet piling. However, there were clear differences in the microbial communities in the corrosion layers of tubercles, which were also different to the microbes present in adjacent seawater. The overall results suggest that the presence of orange tubercles, a single measurement of water quality, or the detection of certain general types of microbes (e.g. sulfate reducing bacteria) should not be taken alone as definitive indications of accelerated corrosion.

## INTRODUCTION

Microbiologically/microbially influenced corrosion (MIC) refers to the deterioration of materials due to the presence of microbes and/or biofilms attached to the surface (1). A special case of MIC is accelerated low water corrosion (ALWC), which is associated with the rapid corrosion of metallic structures at the low tide water level (Fig. 1) (2). The corrosion rates due to ALWC can be up to several mm/yr, compared to around 0.05 mm/yr which is typical for steel in seawater in non-ALWC situations (3). Untreated ALWC can lead to significant degradation of steel support structures and subsequent loss of structural integrity. ALWC is often indicated by orange tubercles that form on the surface of the steel at the low tide water level. The tubercles consist of a combination of corrosion products and complex microbial biofilms including microbes deemed responsible for the corrosion. Beneath the visible orange surface of the tubercle, a black layer is commonly observed and when removed exposes a bright shiny steel surface with localised pitting indicating corrosion. By studying the microbes present in the orange tubercles associated with ALWC it is hoped that a better understanding of the mechanisms involved, and hence improvements in detection and corrosion prevention, can be made.

**FIG 1.**
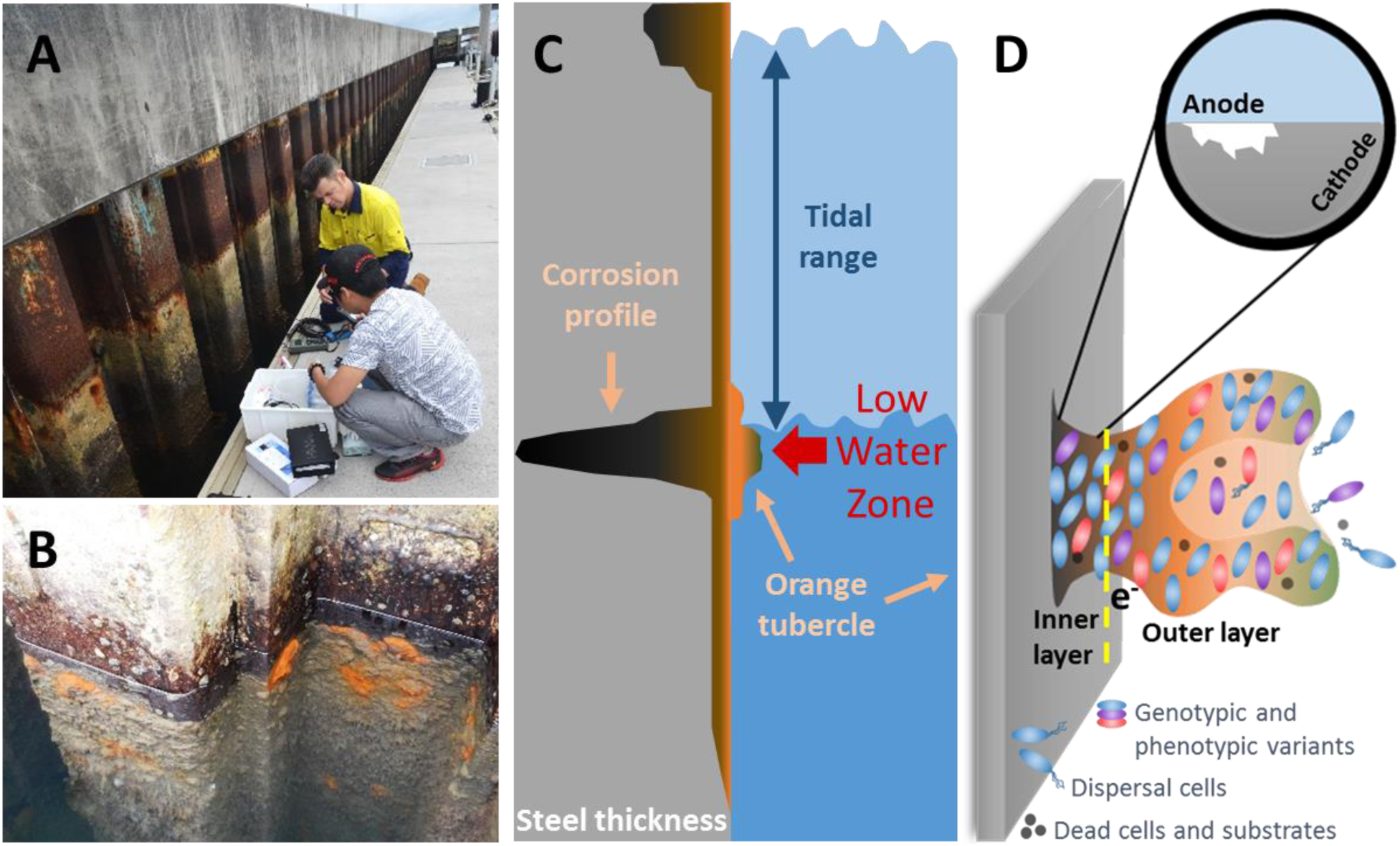
(A) Steel sheet piling being inspected, (B) orange tubercles on steel sheet piling just below the water level, (C) schematic diagram of ALWC and (D) schematic diagram of microbes and layers of orange tubercles.

Biofilms of cells from various microbial species and their extracellular polymeric substances can form on metal surfaces that can subsequently lead to MIC via a range of processes. The exact corrosion mechanisms involved in MIC are likely to depend on the microbes present and environmental conditions. Changes in the chemistry at the metal/solution interface due to metabolic by-products and the direct withdrawal of electrons from the metal are the two most commonly proposed degradation mechanisms (4–6). As the microbial communities involved in MIC are complex and can change spatiotemporally as the organisms adapt to survive in habitats with fluctuating substrate availabilities, it is likely that multiple processes take place that lead to the rapid corrosion associated with MIC. For example, sulfate reduction and sulfide oxidation (parts of the sulfur cycle) are hypothesised to be key mechanisms in accelerated low water corrosion (7, 8).

A wide variety of microbes exist in marine environments, many of which can colonise surfaces to form biofilms (9). ALWC occurs on surfaces at the low tide level where the biofilm is predominantly immersed, but also has periods of exposure above the water level. A small number of studies have investigated the microbial communities present on steel structures in relation to ALWC. One such study (7) found differences between the inner (dominated by sulfate reducing bacteria, SRB) and outer layers (dominated by sulfur oxidising bacteria, SOB) of ALWC tubercles, while another reported a diverse community of bacteria and archaea with a dominant group of methanogenic archaea (10). It is important to note that while they are often linked as being key to MIC, SRB have been found at both sites with normal (i.e. not accelerated) low water corrosion as well as those with ALWC. However, a higher diversity of SRB as well as specific genera of SRB have been reported to be present at ALWC locations compared to normal low water corrosion sites (11). There have also been a number of previous attempts to isolate microbes from ALWC tubercles for use in additional (e.g. corrosion) studies, although typically the microbes isolated have not correlated well with the dominant microbes found in community analysis (7, 12, 13).

One of the other factors that potentially affects the likelihood and location of ALWC is the metallurgy of the sheet piling. Several reports (2) indicate a preference for ALWC to occur in the centre of the out pans of U-profile sheet piling. There are also suggestions that older sheet piling may be more susceptible to ALWC (14). R. E. Melchers et al. (14) have proposed that both of these effects may be due to differences in steel segregation and composition at the centre of the steel sample for the older steel types. However, others report that steel type, and by extension steel composition, did not seem to be a factor for ALWC (2). At least one major steel sheet pile manufacturer markets a sheet piling steel grade with a chemical composition that apparently has been specifically designed to reduce ALWC (15).

In this study, we performed a detailed characterisation of the microbial communities present in orange tubercles at the low tide level on two different types of adjacent steel sheet piling. The overall hypothesis of this work was that the microbial communities present in different layers of an orange tubercle, the nearby seawater and on different areas of the steel structure (with/without orange tubercle, for different steel types, with/without corrosion attack underneath tubercles) would be significantly different. To examine this, we carried out 16S rRNA gene metabarcoding of biomass from the inner and outer layers of numerous tubercles, steel surfaces without tubercles present and for the nearby seawater. One orange tubercle sample, with a pitting interface, was also used to isolate bacteria, which were identified by Sanger sequencing their 16S rRNA genes and phylogenetic analysis. Metallurgical studies of the two different steel types characterised the composition and microstructure of the steels.

## RESULTS

### Seawater

The values of selected measured environmental parameters of the seawater from close proximity to the steel sheet piling, are provided in Table S1, which are reasonably typical for seawater. Of specific interest to ALWC are the dissolved inorganic nitrogen levels. The concentration of ammonia was very low (0.02 mg/L) and the nitrate and nitrite levels were below the levels of reporting (<0.02 mg/L).

### Steel

The elemental composition of the two different steel types was found to be within the standard ranges used in structural grade carbon steels (Fig. S1). For the majority of the elements, the differences in percentages between the old and new sheet piling steels were relatively minor, apart from the copper levels, where the old steel type only had 0.01 wt.% while the new type had 0.33 wt.%.

Examples of the microstructure of the two steel types are given in Fig. 2. The grain sizes of the old steel type (mean intercept length ∼13 μm) were larger than the grain sizes of the new steel type (mean intercept length ∼7 μm). Both steel types had had a mixed pearlite/ferrite microstructure, with obvious banding in the old steel type. The ferrite-pearlite ratio of the old steel and the new steel were found to be 33 to 67% and 23 to 77% respectively. An increased level of inclusions was observed for the old steel type (∼0.7%) compared to the new steel (0.4%). Measurements for each of the aforementioned microstructural parameters were made through the cross-section of each of the samples and no obvious variations were found as a function of measurement location.

**FIG 2.**
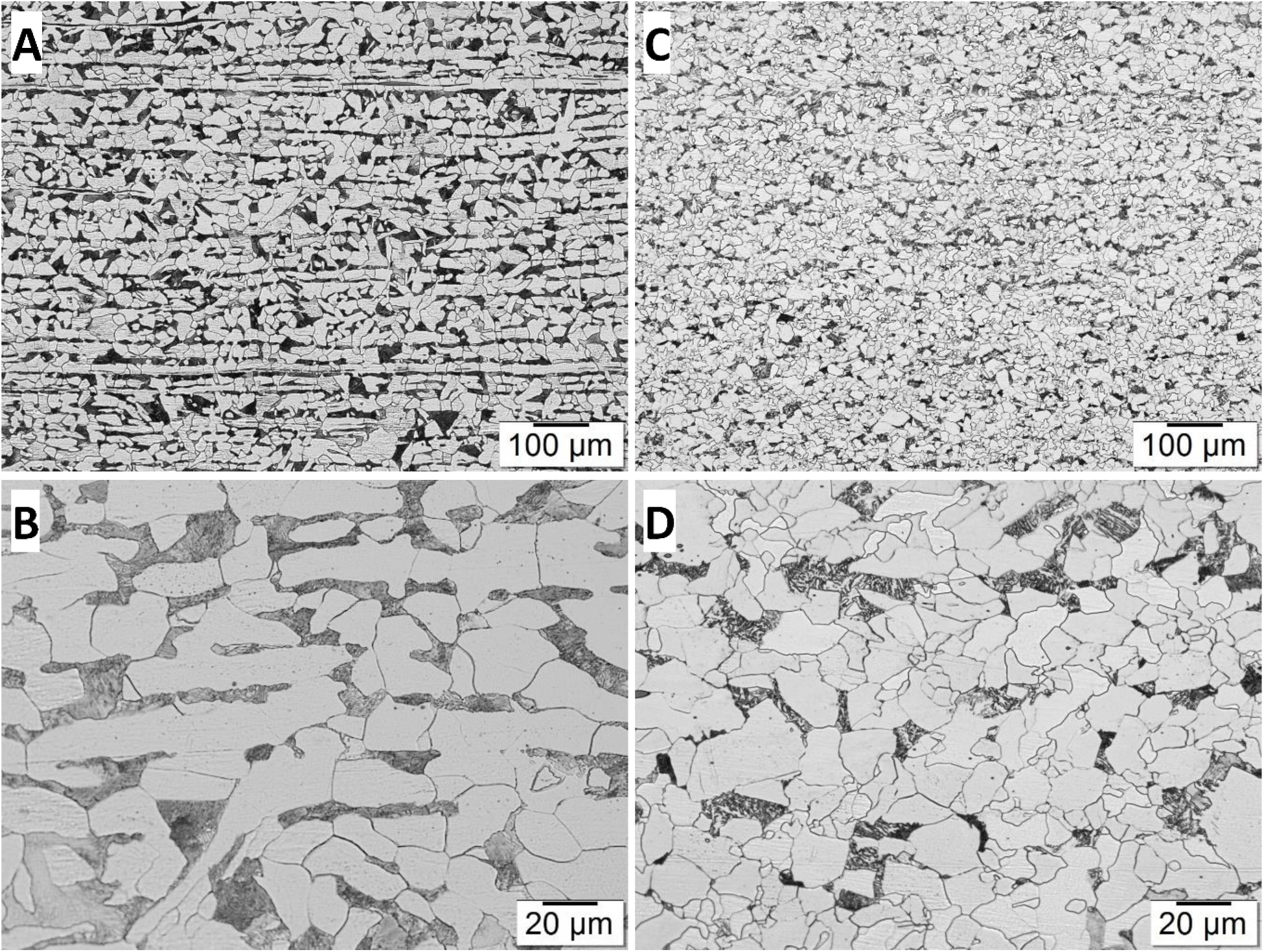
Microstructure of the two sheet piling steel types, from Australian Iron and Steel, Pt Kembla (old, A and B) and ArcelorMittal, Luxembourg (new, C and D), at different magnifications, pearlite phase (black grains) and ferrite phase (white grains).

### Pitting/Localised corrosion

Pitting, i.e. localised corrosion of the metal surface, was observed beneath certain orange tubercles (e.g. see Figs. 3 and S2) and in some cases was estimated to be up to a few cm in depth (Fig. S3). There was extensive pitting present under all tubercles removed from the old steel type (Fig. S3 A–D, G and H), whereas the new steel type was relatively smooth and devoid of cavities (Fig. S3 E and F).

**FIG 3.**
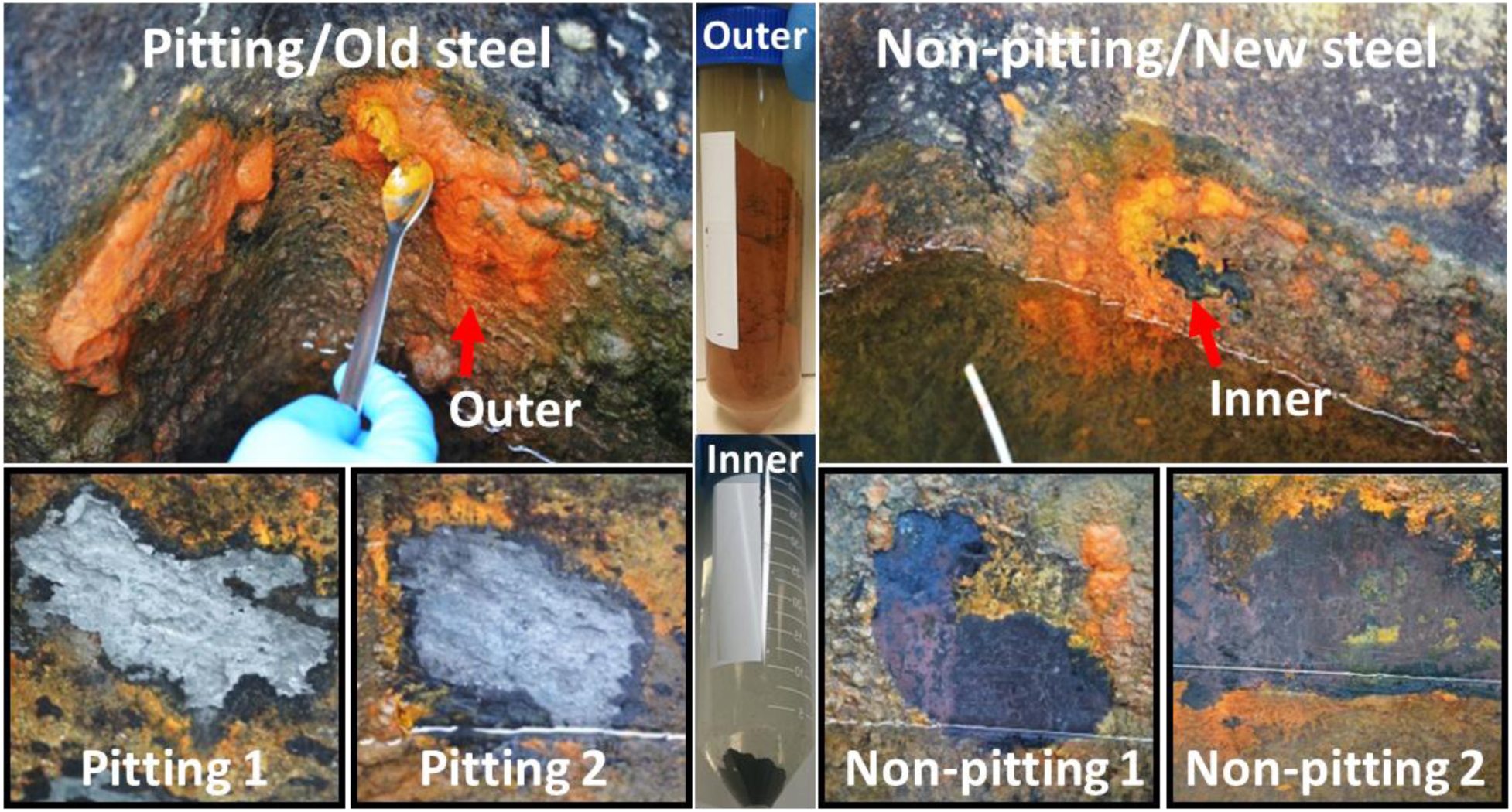
Photographs of examples of orange tubercles (top) and cleaned steel surfaces (bottom) for the two different steel sheet piling types, (left; pitting/old steel) Australian Iron and Steel (Port Kembla) steel sheet piling and (right; non-pitting/new steel) ArcelorMittal (Luxembourg) steel sheet piling.

### Microbial communities – metabarcoding

The phylum level microbial communities of the various samples investigated are presented in Fig. 4A. The most dominant phylum was Proteobacteria, which made up between 47.3 and 86.2% of the microbes detected in individual samples. Bacteroidetes were present at reasonable levels in most of the samples tested, being the highest in the seawater (29.6%) and the lowest (1.1%) in the non-pitting inner 2 sample. The archaeal phyla detected included Euryarchaeota and Thaumarchaeota, but they were recorded at relatively low levels across the different orange tubercle samples, ranging from undetected to ∼20%, while none were present in the seawater. Actinobacteria and Cyanobacteria comprised 4.4% and 6.3%, respectively in the seawater, but were <1% in all but one of the tubercle samples. The combination of Unassigned and Others (phyla detected at levels <0.5%) comprised between 8.9% and 36.5% of the detected microbes in individual samples.

**FIG 4.**
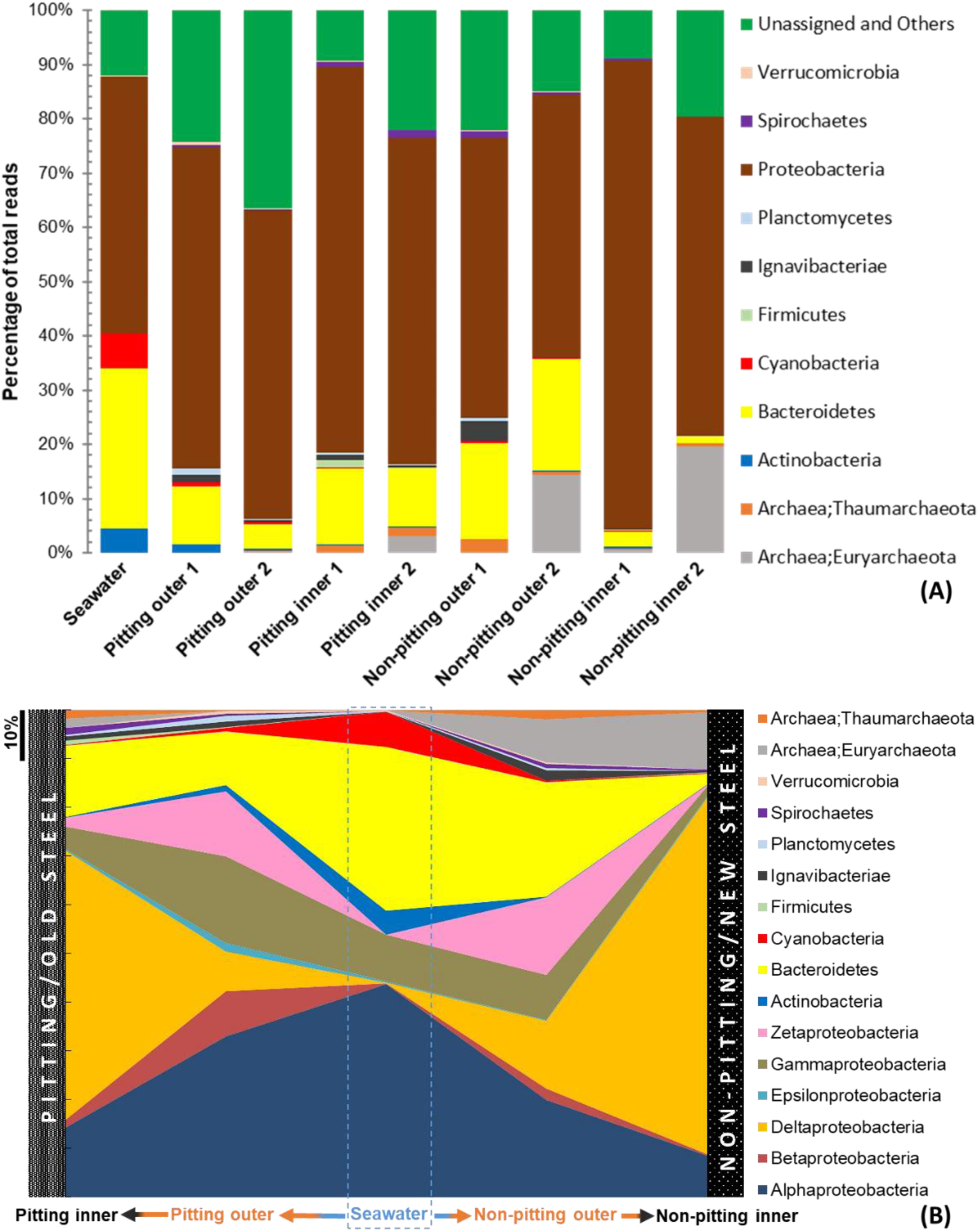
(A) Microbial communities of different samples at the phylum level as determined by metabarcoding. Others, including phyla with levels identified to be <0.5% in all samples and unassigned phyla, were grouped into “Unassigned and Others”, and (B) Major phylum/class bacterial groups plotted for the different sample locations (“Unassigned and Others” removed). The data from the two test locations for each steel type have been averaged to help simplify presentation.

To illustrate the distribution of microbial communities present in the different regions of the orange tubercles as a function of steel type, the results obtained for individual tubercles tested for the two steel types were averaged and plotted in Fig. 4B, together with the seawater results for comparison. For this figure the Proteobacteria were expanded to the class level, while the Unassigned and Others (here defined as phyla detected at levels <0.5%) were removed from Fig. 4B. Alphaproteobacteria were more dominant in the seawater (38.5%), with the relative proportions lower in the outer tubercle layer and then further reducing in the inner layer closest to the steel (∼10%). Similar trends of reducing abundance from seawater through to the inner layer of the tubercles were observed for the Actinobacteria, Bacterioidetes and Cyanobacteria. In general, the outer layers of the tubercles had increased levels of Betaproteobacteria and Zetaproteobacteria, while the inner layers tended to be dominated by Deltaproteobacteria (∼50%). Gammaproteobacteria showed slight changes in proportion across the seawater and outer samples (∼10%) with a decrease in the inner layers (∼2%).

The within-sample rarefaction (Alpha diversity), as determined by QIIME using the OTU table, is shown as the basis of phylogenetic diversity whole tree and Shannon’s metrics, and plotted in Fig. S4A. The sequencing depth was considered to be good for each sample as it reached the samples’ richness. The rarefied OTU table was run by a matrix of distances among all samples, which is illustrated in the principal coordinates (Fig. S4B). A comparison of the community compositions was performed with 1,000 permutations using the Bray-Curtis metric. Significant differences (*R* > 0.7, *p* < 0.05) were shown by ANOSIM *R*-statistic in samples (i.e., inner, outer or seawater) and sources (i.e., pitting inner/outer, non-pitting inner/outer or seawater); but not significant in terms of the type of steel on which the tubercles were located (i.e. pitting/old steel type or non-pitting/new steel).

A heatmap produced at the microbial family level (including 77 microbial families) was integrated with a clustering dendrogram using Pearson’s correlation and built in METAGENassist. The results showed some distinct groupings occurred for the different sample locations. For example, the Desulfobacteraceae and Desulfovibrionaceae were present in the inner layers of the orange tubercles but there was a negative correlation for these families in both the tubercle outer layers and in the seawater. Further details are provided in Fig. S5 in the supplementary material.

To provide information at the genus level, the taxa present in the five sample locations at levels ≥1% are plotted in Fig. 5. Specific taxa that were consistently identified at the species levels across all of the 5 sample locations are provided on the right hand side of the bars (Fig. 5). All taxa present at levels <1% were grouped together and plotted as a single group called minor taxa, to show their relative abundance. In general, the non-pitting inner and non-pitting outer samples had less diverse microbial compositions than those of pitting inner and pitting outer samples, respectively.

**FIG 5.**
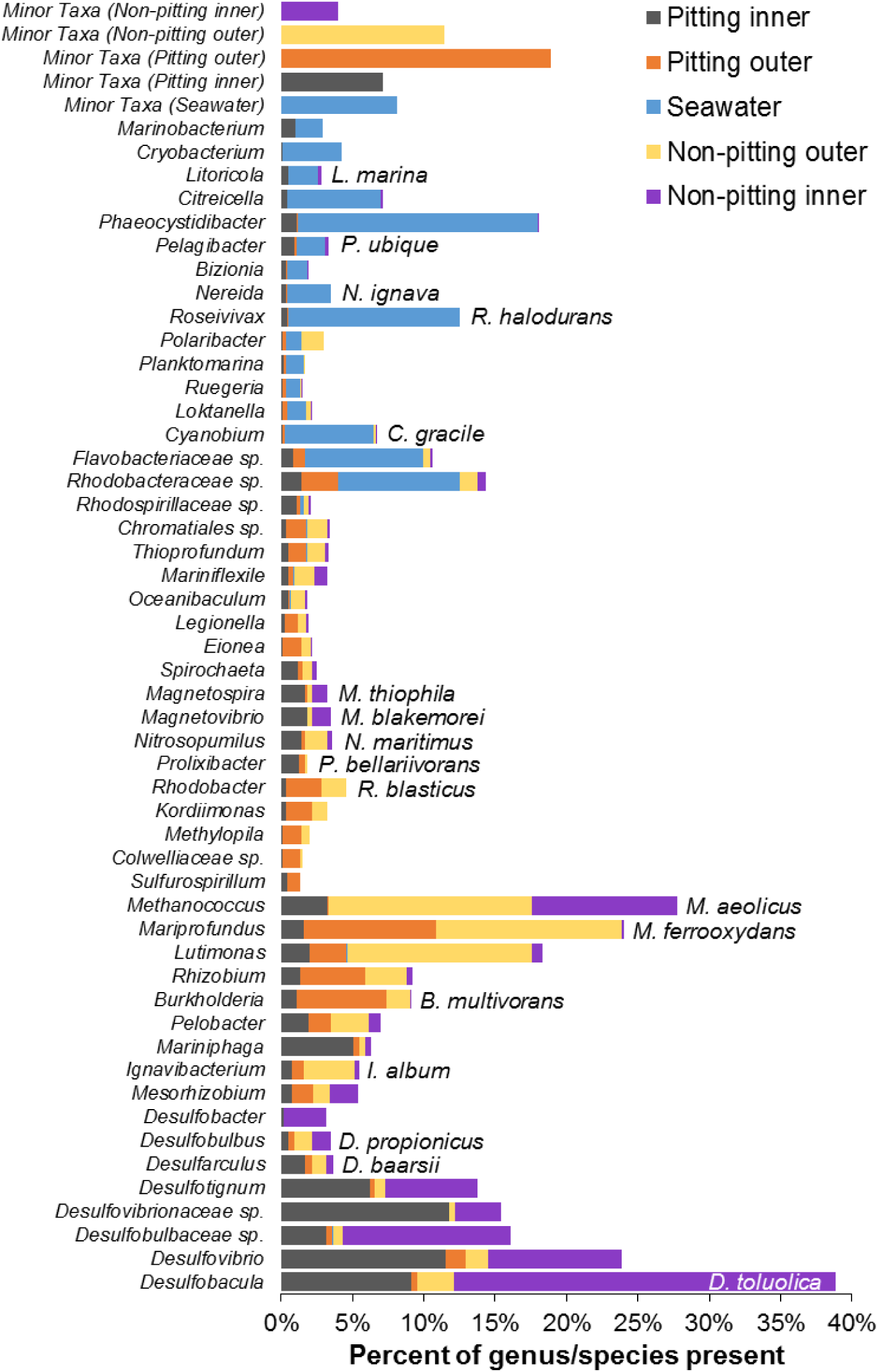
Percentages of families and/or genera/species present in the various sampling locations and layers (see code in figure) according to metabarcoding. Note that pitting occurred below orange tubercles on the old type steel, while the new type steel did not show pitting beneath orange tubercles.

The most dominant genera present in the seawater sample were *Phaeocystidibacter*, *Roseivivax halodurans*, and species of the families Flavobacteriaceae and Rhodobacteraceae, while the outer layers of the tubercles had high levels of *Methanococcus aeolicus* (an archaean), *Mariprofundus ferrooxydans*, *Burkholderia multivorans* and *Lutimonas* (Fig. 5). The inner layers of the tubercles of both steel types contained a range of Deltaproteobacteria (lower eight taxa in Fig. 5) with relatively high proportions of *Desulfobacula toluolica*, *Desulfovibrio* sp., *Desulfotignum* sp. and a species of each of families Desulfobulbaceae and Desulfovibrionaceae.

In comparing microbes from the inner tubercle layers on both steel types (old and new), *D. toluolica* and species of Desulfobulbaceae were less abundant on the old steel type than on the new. Conversely, more Desulfovibrionaceae species and *Mariniphaga* (Bacteroidetes) were in the inner layers of the old steel type compared to the new steel. Rhodobacteraceae species were in higher proportions on the old steel compared to new, and *Prolixibacter bellariivorans* and *Sulfurospirillum* were only present on the old steel. *B. multivorans* were clearly more abundant on the old steel, while *M. aeolicus* were present at greater levels on the new steel.

METAGENassist was used to provide the putative phenotypes of the microbes identified by metabarcoding and a summary is presented in Fig. 6. However, the functions of the majority of microbes (∼60%) from each sample were unknown. Overall the six most dominant traits identified were sulfate reduction, ammonia oxidation, dehalogenation, nitrite reduction, nitrogen fixation and sulfide oxidation. Generally higher metabolic function percentages were in the pitting rather than in non-pitting tubercles and more in the inner layers compared to outer layers for tubercles on both steel types. Particularly sulfate reduction, was greater in the seawater and inner layers than the outer layers of the tubercles, ammonia oxidation was lowest in non-pitting tubercles and there was little difference in sulfide oxidisation among corrosion layers. Across the samples, dehalogenation and nitrogen fixation were highest in the pitting inner layer.

**FIG 6.**
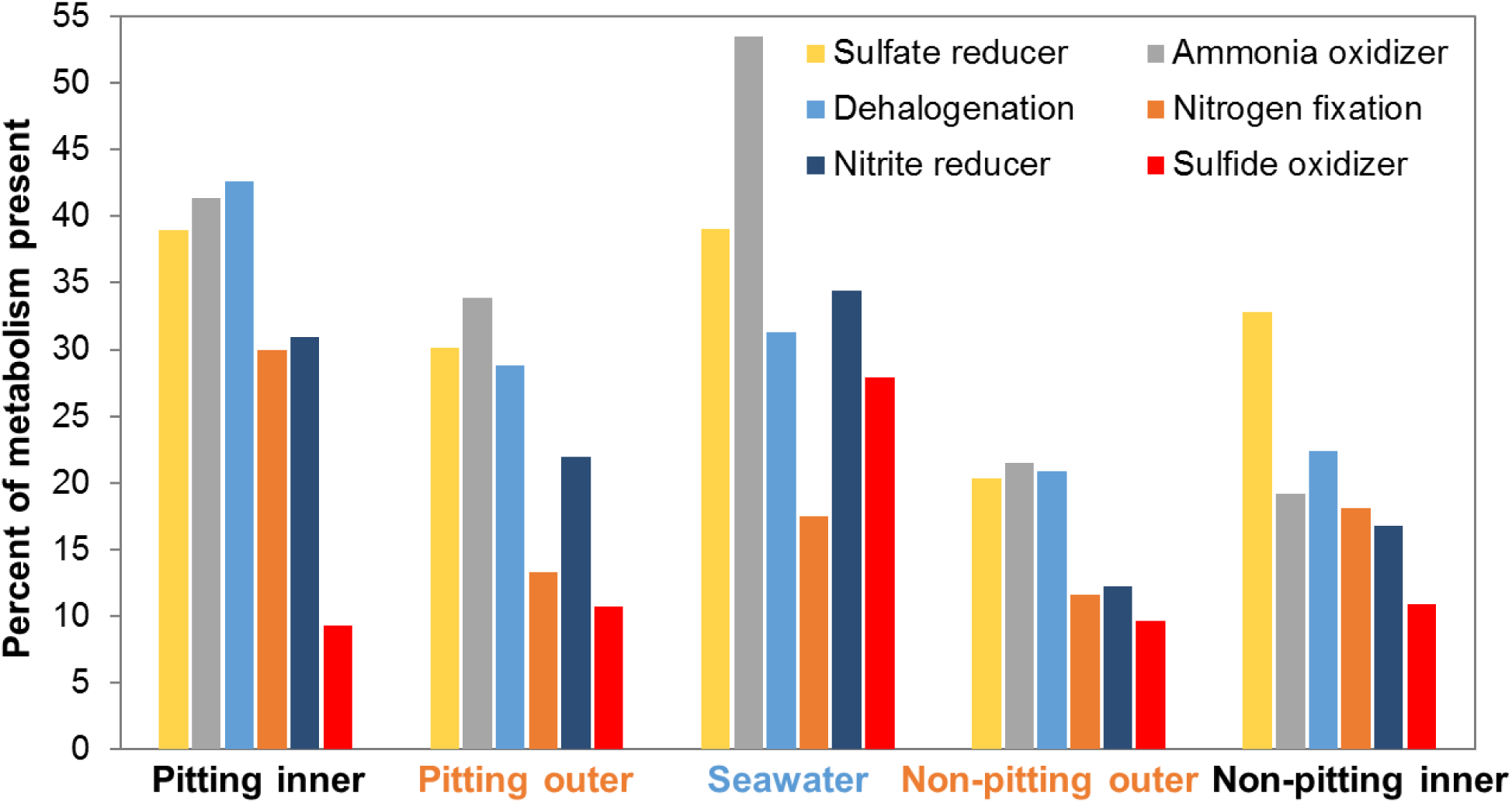
Putative functions of OTUs obtained for the various test locations, as analysed by METAGENassist.

### Microbial communities – pure cultures

A total of 119 pure culture bacterial isolates were obtained from a combination of the inner and outer layers of an orange tubercle from one of the pitting/old steel locations. These isolates produce a culture collection which can be used in subsequent laboratory-based corrosion tests. The phylogeny of the isolates is shown in a circular phylogenetic tree in Fig. 7 including details of close relative identities. Further details, including the relative abundance percentages of each genus based on plate counts, are provided in the supplementary material (Table S2). For both the aerobic and anaerobic culturing conditions, a more diverse bacterial population was able to be isolated from the outer layer and it also had a higher total relative abundance of bacteria (according to plate counts) compared to the inner layer. *Bacillus* was the most dominant genus of bacteria isolated aerobically, while *Mariniphaga* and *Prolixibacter* (all were *P. bellariivorans*) and *Clostridium* were the most common genera isolated anaerobically. The different culturing conditions (medium composition and atmosphere) would have influenced the isolates acquired. The majority (∼95%) of the isolates could grow on either MA or R2A-SS, while one isolate each of *Sphaerochaeta* and *Yeosuana* (an aerobe) were only isolated on THA^+^. In aerobic cultivation, MA and R2A-SS collectively facilitated a greater diversity of microbes compared to other media. The addition of lactate and ferrous ammonium sulfate to growth media (TSA^+^ and MA^+^) facilitated the isolation of fewer bacteria which could be due to the media being more selective.

**FIG 7.**
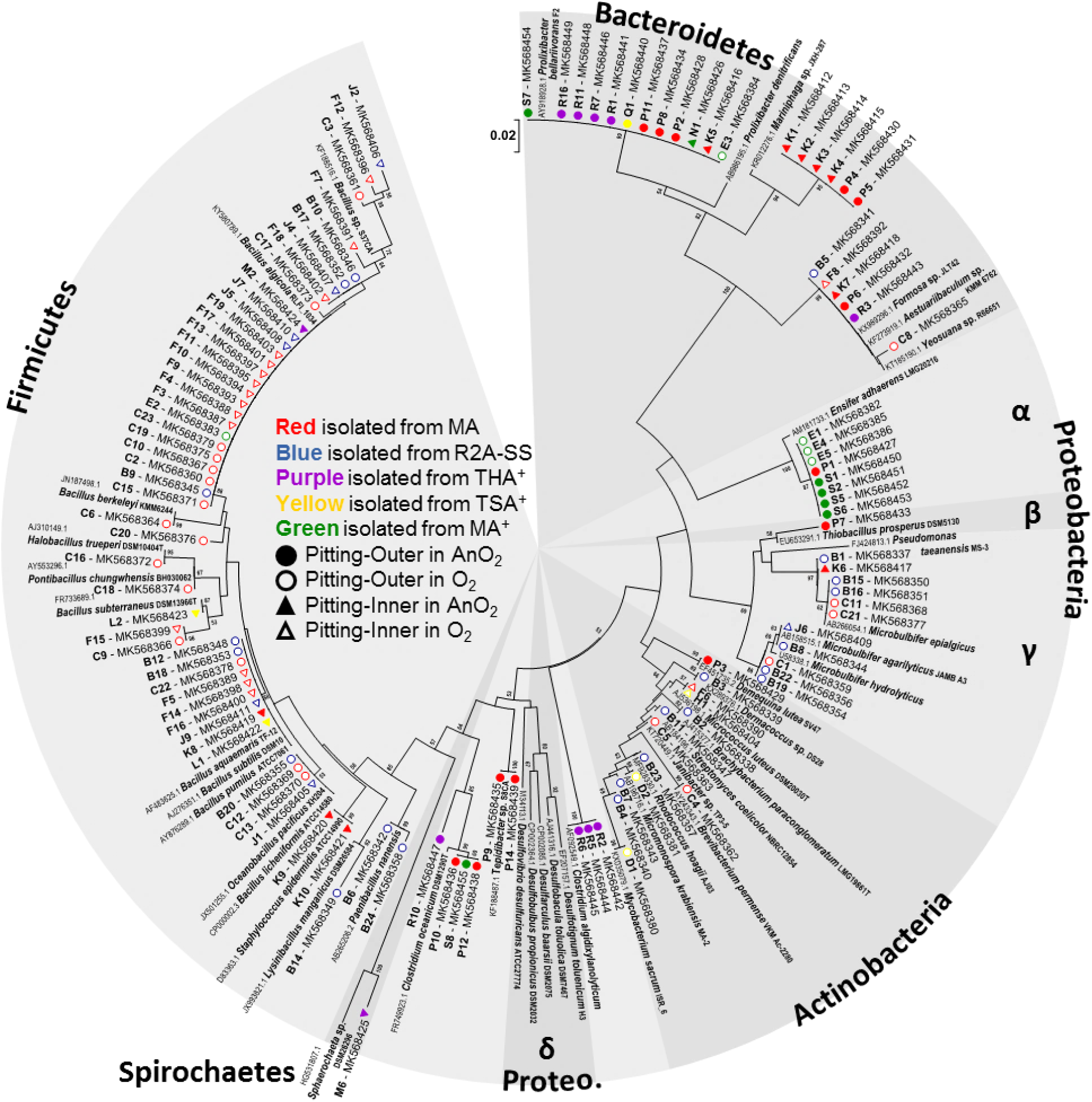
Neighbor-Joining phylogenetic tree created with MEGA7 for 16S rRNA sequences of bacteria isolated from tubercle layers on a pitting old steel. Evolutionary distances were analysed using the bootstrap test from 2000 replicates. GenBank accession numbers are noted after isolate codes and before names of reference sequences.

## DISCUSSION

### Physico-chemical properties of seawater

The major water parameters analysed in this study were within the ranges of those reported in a previous study examining the water quality of region close to the area we studied (16). Others have proposed a link between the concentration of dissolved inorganic nitrogen (DIN) present in seawater and likelihood of ALWC (17). Our, albeit one off, measured DIN levels were quite low (0.02 mg/L), which based on the aforementioned theory would suggest a reduced likelihood of ALWC. However, we clearly observed pitting, which have directly correlated with corrosion (Fig. 3) underneath orange tubercles on the old steel type.

### Steel microstructure

We found a clear difference in the extent of pitting (i.e. localised corrosion) beneath orange tubercles for two different steel types that are located adjacent to each other and installed at the same time. Given that we saw little difference in the microbial communities present in the tubercles on the two different steel types this suggests that something to do with the steel properties has possibly influenced the corrosion process. Previous work has shown that banding of the steel microstructure (18) and MnS inclusions (19) can be the cause of increased levels of localised corrosion of steel. Banding and increased levels of inclusions were observed for the old grade of steel which had increased corrosion. Another factor which may potentially affect the likelihood of ALWC is the steel composition, although there are differing reports on this (2, 15, 20). While some differences were found in the composition of the two steels investigated (e.g. increased levels of copper in the steel without the pitting) they are relatively small, and the potential of copper alone to reduce ALWC is not clear (20). Given the number of variables in play, further work is required to determine which, if any, of these material properties are responsible for the increased levels of corrosion in the old steel type.

### Microbes – metabarcoding and METAGENassist

The tubercles formed had clear differences in the microbial communities in the different layers (regardless of steel type), which were also clearly different to the microbes present in adjacent seawater in this study. The bacterial compositions determined by metabarcoding at the phylum level in this study were similar to those in a previous study, in which the rust layers from steel plates immersed in coastal seawater for 6 months and 8 years were examined (21). In particular in the latter samples (steel exposed for 8 years in seawater) and in ours (steel immersed for 7 years in seawater), Proteobacteria and Bacteroidetes were the abundant phyla of bacteria (Fig. 4); the Firmicutes however were higher in samples from X. Li et al. (21) compared to our samples.

An earlier study of the microbes on seawater immersed steel coupons (22) generated results that concurred with our findings of relatively few archaea compared to bacteria overall (Fig. 4A), and bacterial communities clustered by tubercle location (inner and outer) and water (Fig. 4B). Alphaproteobacteria and Gammaproteobacteria distribution (with abundance changing from high in seawater through to the lowest in tubercle inner layer) concurred between the studies. Some differences, however, were also observed with Bacteroidetes distributions, for example, being high in seawater through to lowest in the tubercle inner layer in our study (Fig. 4), but the opposite being reported in (22).

There were some similarities between our bacterial findings and a third MIC study (Zhang 2019), particularly with similar abundances of Proteobacteria and Bacteroidetes. In general, other studies have reported more Firmicutes (X. Li et al. (21), S. Celikkol-Aydin et al. (22) – mostly *Defulfotomaculum*) and sometimes more Actinobacteria (22, 23), compared to our results, but the Proteobacteria and Bacteroidetes are abundant phyla in all these studies.

There are several potential reasons for the variations in microbial communities in corrosion samples, including different local environmental conditions (e.g. nutrients, temperature, other water compounds), water flux (24) and immersion times, which influence specific marine microbial communities (21, 25). Less diverse microbial communities have been reported with increasing immersion time (6 months versus 8 years; X. Li et al. (21)). The structures studied in our work had been immersed for ∼7 years prior to sampling and the microbial diversity was similar to that in the 8 year sample previously reported (21).

It has been suggested (26) that algae then cyanobacteria are primary terrestrial biofilm colonisers and data from S. Celikkol-Aydin et al. (22) add credence to the notion they could be primary colonisers in marine metallic corrosion settings. We did not specifically explore algae, and our cyanobacteria were largely only in the seawater sample. Our tubercles were likely mature biofilms (the steel was in place for ∼7 years) making initial colonisers unfeasible to comment on.

Several microbial phenotypic properties were suggested from the METAGENassist analysis. Of particular note are aspects of sulfur (sulfate reduction and sulfide oxidation) and nitrogen (nitrogen fixation, ammonium oxidation and nitrite reduction) cycling in pitting and non-pitting samples from the inner and outer of the tubercles (Fig. 6). The seawater microbes were enriched for many of these phenotypes and often more so than the tubercle samples – e.g. sulfate reduction and ammonium oxidation. Sulfur cycling, especially the sulfate reduction component, is well known with respect to MIC and indeed is frequently deemed a critical phenotype for this type of corrosion. METAGENassist showed a trend for more sulfate reducers in inner regions of both pitting and non-pitting tubercles compared to respective outer regions (Fig. 6). This phenotype was high in the seawater and the likely source of ALWC/MIC sulfate reducers as our previous research could not confirm their source to be adjacent sediments (12). METAGENassist created heat map shows a positive correlation for putative sulfate reducers (Desulfovibrionaceae and Desulfobacteraceae) in inner samples compared to outer and seawater samples (Fig. S5).

Sulfate reducing prokaryotes have been widely implicated and studied in ALWC/MIC (5). *Desulfovibrio* sp. have previously been suggested to be a main cause of corrosion by the direct electron transfer mechanism (27), and both our pitting and non-pitting inner samples contained this genus, other Desulfovibrionaceae and other known sulfate reducers (*Desulfarculus baarsii*, *Desulfotignum* sp., Desulfobulbaceae and *Desulfobacula toluolica*) (Fig. 5). METAGENassist supports the hypothesis that sulfate reduction is a potential microbial phenotype in the inner zones of our collected tubercules. According to metabarcoding, our non-pitting inner samples had more sulfate reducers compared to the pitting samples (Fig. 5) but METAGENassist showed slightly more sulfate reduction in pitting samples compared to non-pitting samples (Fig. 6). As such, in this work corrosion could not be directly correlated with putative sulfate reducers as determined by both metabarcoding or METAGENassist. This finding is at odds with the general status in the ALWC/MIC field of a direct relationship between sulfate reduction and corrosion, and therefore warrants more research to provide clarity.

The sulfur cycle could have involved the sulfur respirer *Pelobacter* (Desulfuromonadaceae), which are of marine origin (28) and were found in all inner and outer samples (Fig. 5). The sulfur oxidisers *Thioprofundum* sp. (Gammaproteobacteria) and *Sulfurospirillum* sp. (Epsilonproteobacteria) were largely in outer tubercle samples compared to inner or seawater (Fig. 5). High sulfide oxidation was found in the seawater according to METAGENassist (Fig. 6) but since we could not identify seawater bacteria with this phenotype, this warrants further investigation.

The most abundant archaeon detected was the marine methanogen *Methanococcus aeolicus* (Euryarchaeota) in the non-pitting inner and outer locations, and in lower abundance in the pitting inner sample (Fig. 5). *Methanococcus* and *Methanothermococcus* were the abundant archaea detected in rust samples on steel surfaces immersed in seawater for 2.5 yr. There was an inverse relationship between these archaeal genera and location in the rust, with high *Methanococcus* in the outer and high *Methanothermococcus* in the inner (23). The marine ammonium oxidiser *Nitrosopumulis maritimus* (Thaumarchaeota) was present in low abundance in inner and outer samples. These have been reported from MIC settings (23) but their role in corrosion in non-oil pipeline locales is yet to be explored. Their phenotypes of methanogenesis and ammonium oxidation are possibly important to corrosion processes.

### Microbes – pure cultures

All of the media used in this work to isolate pure cultures were generally non-selective, but they varied in their nutrient (R2A is a low nutrient medium relative to the others) and salt levels (all but TSA^+^ were prepared to seawater salinity with RSS but TSA^+^ has 0.5% NaCl). We used the seawater salinity isolation media so that the broadest spectrum of marine organisms could be obtained. The supplements to TSA, THA and MA were used to enhance the isolation of anaerobes and, in particular, TSA^+^ for sulfate reducers. This could have provided a selective aspect since substantially less bacteria grew on MA^+^ compared to MA. Our specific use of TSA^+^ as the medium to facilitate isolation of SRB was unsuccessful. Indeed, we isolated no SRB or facultative SRB (e.g. *Vibrio* spp.) from any medium despite good proportions of potential SRBs (Deltaproteobacteria) being detected, particularly in the pitting inner samples, from metabarcoding. I. Lanneluc et al. (13) using similar methods as ours with 8-week immersed metal coupon biofilms also could not isolate SRB. These bacteria are considered to play a main role in corrosion (5) and the isolation of corrosion associated SRB is a high priority to allow use in future MIC/ALWC studies.

We isolated bacteria from five phyla (in abundance order): Firmicutes (many *Bacillus* spp.), Proteobacteria (mostly *Ensifer adhaerens* (Alphaproteobacteria) and three different *Microbulbifer* spp. (Gammaproteobacteria)), Bacteroidetes (mostly *Prolixibacter bellariivorans* and *Mariniphaga* sp.), Actinobacteria (at least nine different genera and species), and Spirochaetes (*Sphaerochaeta* sp.). This is a similar diversity of pure cultured bacteria reported from rust samples on steel that had been immersed in seawater for 8 years (25) and in seawater for up to 8 weeks (13). Our former four phyla are consistent with those reported by X. Li et al. (25) and I. Lanneluc et al. (13), but one isolate, *Sphaerochaeta* sp. is unique to our work, novel for corrosion environments and justified the use of THA^+^ medium from which our *Sphaerochaeta* sp. was isolated. Most *Sphaerochaeta* sp. are reported to come from freshwater or gut ecosystems but the anaerobic, psychrophilic, marine *Sphaerochaeta multiformis* was sourced from subseafloor sediments collected near Japan in the north-western Pacific Ocean (29). However, the closest bacterium to our strain was *S. pleomorpha* (from freshwater sediment) with 89.1% identity and *S. multiformis* was 87.1% identical. Thus, our *Sphaerochaeta* sp. strain is likely a novel species.

Most of our isolates were *Bacillus* sp., specifically *B. algicola* and *B. aquimaris*, despite extremely few sequences from this genus present in our metabarcoding data. It could be that the *Bacillis* spp. were present in our corrosion samples as spores, which could have eluded lysis in our protocol, thus leading to few sequences from them in metabarcoding. Others have also isolated *Bacillus* spp. in abundance from corrosion samples. I. Lanneluc et al. (13) studied metal coupons immersed in seawater for 8 weeks and used MA to find that 38% of isolates were Firmicutes (Class Bacilli), most of which were *Bacillus* spp. More relevant to our work, due to a similar marine water immersion time of 7–8 years, X. Li et al. (25) found *Bacillus* spp. abundant isolated organisms, but not present in their metabarcoding data (30). *Bacillus* spp. have been reported to have different roles in corrosion, ranging from promoting to inhibiting corrosion. *Bacillus* spp. can produce extracellular polymeric materials (EPS) which bind metal ions leading to anodic dissolution and promote corrosion (31). Conversely, *Bacillus brevis* inhibit corrosion by producing antibiotics against SRB and iron oxidising bacteria (32). The true abundance of *Bacillus* taxa in MIC needs to be clarified and, if found to be abundant, then their specific roles need to be clarified.

*Microbulbifer* spp. are known manganese oxidisers (33) and bacteria with this phenotype are associated with corrosion of iron and steel (34). They also could be common surface colonizers contributing to initial MIC biofilm development as reported for several bacterial families including Alteromonadaceae (*Microbulbifer*) (35).

MA^+^ was very successful in isolating our only candidate of Alphaproteobacteria, *Ensifer adhaerens*, which grew both aerobically and anaerobically. *Ensifer* spp. are common surface colonisers (36) and have a symbiotic relationship with *Bacillus* where they can promote *Bacillus* spore germination and receive vital nutrients that are produced during spore activation (37). These and other phenotypic features (e.g. As(III) oxidation (38) and nitrogen fixation (39)) may explain the ubiquity of *E. adhaerens* in our samples.

*Prolixibacter bellariivorans* grew on all media under anaerobic incubation and most were from the pitting outer sample. *Mariniphaga* spp. were mainly isolated anaerobically on MA from both pitting inner and outer samples. Both these genera are in the Prolixibacteraceae family in the Bacteroidetes. *Prolixibacter* have been reported to be anaerobic iron oxidisers using nitrate reduction (40, 41). The role of *Mariniphaga* in ALWC has yet to be determined. All other Bacteroidetes isolates were in the family Flavobacteraceae, which have been reviewed as primary colonisers in MIC (9) and from which isolates were reported from other MIC studies (13, 25).

## MATERIALS AND METHODS

### Sampling

Orange tubercles (∼7 mm thickness and ∼60–100 mm diameter) which had formed at the low tide water level on a steel sheet piling wall in a marine harbour in Port Philip Bay, Victoria, Australia were sampled (Figs. 3, S1**–**S3). The sheet piling studied consisted of two different steel types (one manufactured by Australian Iron and Steel, Port Kembla, Australia (old steel) and the other by ArcelorMittal, Luxembourg (new steel) – see analysis below) of U-piles connected together to form a single ∼80 m straight length. The wall was constructed in 2009, and duplicate tubercles were sampled in 2016 from each of the two different steel types, making a total of four sampling locations.

When the orange tubercles were removed, bright shiny pitted steel was observed beneath the old steel type and samples acquired were called pitting 1 and pitting 2 (Figs. 3, S1–S3). The steel below the new steel type had no pitting (i.e. localised corrosion) and a dark-black appearance and samples acquired from this region were called non-pitting 1 and non-pitting 2 (Figs. 3, S1–S3). Each orange tubercle was sampled in a manner that allowed separate specimens of the outer bright orange, thick layer (called “outer”) and inner black, thin layer (called “inner”) to be obtained (Fig. 3). An ethanol-disinfected spoon (Fig. 3) was used to scrape the samples into 50 mL polycarbonate tubes (Falcon™), which were then filled with seawater. All sample tubes were kept on ice during transportation to the laboratory. Seawater adjacent to the sheet piling was collected into a 20 L plastic container for microbial analysis. Orange tubercles were observed on “in” and “out” pans of the U-pile sheets, with all samples in this work taken from “in pan” areas.

At the laboratory, three aliquots of 500 mL of well-mixed seawater taken from the field were filtered through a 0.22 μm mixed cellulose esters membrane (Merck Millipore) using sterile techniques. The filter was stored at –80°C prior to further testing. Tubercle inner and outer samples were individually homogenised using a paddle mixer (Stomacher) at normal speed for 2 min. The final samples were aliquoted into sterile 15 mL sterile polycarbonate (Falcon™) tubes. One aliquot of each sample was kept at 4°C for DNA extraction undertaken on the following day. The remaining tubes were stored at –80°C.

Physico-chemical measurements were taken on site for temperature, pH (HI98185, Hanna Instruments), conductivity (HI8733, Hanna Instruments) and dissolved oxygen concentration (HI9146, Hanna Instruments). A range of other water quality parameters were determined by a commercial environmental testing laboratory (Eurofins, Melbourne).

### Sheet piling

The steel U-pile sheets from two different sources were studied. One was repurposed from a previous harbour structure, which was manufactured by Australian Iron and Steel (Port Kembla) to an AIS 72 lb/ft standard and ∼21 mm thickness. The other, newer U-pile steel was AU25 (Grade 350) manufactured by ArcelorMittal (Luxembourg) and is ∼14 mm thickness. Samples of each steel type were obtained by cutting ∼100×150 cm sections from the top of the sheet pile, using an angle grinder with a thin fibre disk to minimise heat generation (Fig. S1).

Sub-sections were cut from the steel sections using an abrasive water-jet machine (OMAX-1515 MAXIEM, USA). Inductively coupled plasma atomic emission spectroscopy was used to determine the chemical composition of the steels. Samples cut with the long axis in the same direction as the rolling plane (Fig. S1) were used for microstructural analysis. These samples were first manually ground using silicon carbide papers (grit size 500 and 1200) then fine polished using a sequence 5, 3 and 1 μm diamond-based suspensions. Optical microscope images (BX61, Olympus, USA) were taken at a range of magnifications of various locations on the longitudinal plane and rolling plane surfaces to determine the presence of inclusions. Quantification of the inclusion content was determined using thresholding with ImageJ (42).

To observe the microstructure of the steels the polished metal coupons were etched using 2% Nital solution, a combination of 98 mL absolute ethanol (100%) and 2 mL nitric acid (70%). Microscopic images were then taken of various locations on the longitudinal plane and rolling plane surfaces at various magnifications. These images were then used to determine the grain size, using the intercept method as per the ASTM standard E112 (43), and the percentage of pearlite phase present in the steels, using as per the ASTM standard E562 (44). Visual inspections of the sheet piles indicated similar levels of general corrosion (above the water line) and similar numbers/morphology and extent of orange tubercles on the two different steel types.

### DNA extraction, amplification and sequencing of microbial communities

Genomic DNA was extracted from the eight collected samples (i.e. inner and outer layers of tubercles at the four locations) and the seawater filter membrane using the UltraClean™ Microbial DNA Isolation Kit (MoBio, Qiagen). An additional step of heat treatment (65°C for 15 min) following the bead beating step was included in attempts to increase the DNA yield of environmental samples and DNA was quantified using Nanodrop Spectrophotometer, ThermoFisher. Extracted DNA was amplified by PCR using primers (italic parts) 515F (5’-TCG TCG GCA GCG TCA GAT GTG TAT AAG AGA CAG TAT GGT AAT TGT---*GTG YCA GCM GCC GCG GTA A*-3’) and 806R (5’-GTC TCG TGG GCT CGG AGA TGT GTA TAA GAG ACA GAG TCA GTC AGC C---*GGA CTA CNV GGG TWT CTA AT*-3’) for the variable region 4 (V4) of 16S rRNA genes (45, 46) and including the metabarcoding adapters, pads and linker nucleotides. Each reaction tube contained 10-25 ng DNA, 12.5 µL of MangoMix (Bioline), 1 µL of each primer (10 µM), and sterile MilliQ water up to 25 µL. DNA was amplified in a MyCycler^TM^ Thermal Cycler (Biorad) with the following temperature profile: an initial denaturation at 94°C for 5 min; 35 cycles of denaturation (94°C for 45 sec), annealing (55°C for 45 sec), extension (72°C for 45 sec); and a final extension at 72°C for 5 min. Duplicate PCR products of the appropriate length (ca. 382 bp) of each template were pooled to a total volume of 25 µL and transported on dry ice to the Ramaciotti Centre for Genomics (University of New South Wales, Australia) for sequencing using an Illumina MiSeq sequencer.

### Statistical analysis of metabarcoding data

The QIIME version 1.9.1 environment was installed on Australia’s National eResearch Collaboration Tools and Resources cloud-based supercomputing platform. The 16S rRNA gene libraries were filtered using Trimmomatic 0.36 to eliminate low quality sequence reads. PandaSeq 0.19.2 was the tool to join paired-ended reads and discard the joined sequences of less than 250 nucleotides in length. USEARCH 6.1 was used for chimera detection and OTU (Operational Taxonomic Unit) picking. Default identity threshold of USEARCH 6.1 was 97%. The nonchimeric sequences per sample were counted and created the OTU table. Taxonomy assignment was clustered against the 16SMicrobial database from NCBI (downloaded on 29 Mar 2017 and processed by the NCBI 2.2.22 software package). The Alpha diversity was performed for richness quality of the observed OTUs within each sample. Additionally, the Beta diversity was analysed under the algorithm of Bray Curtis and ANOSIM to present the OTU composition among groups of samples. Additionally, the final OTU table and its metadata were input to the METAGENassist (www.metagenassist.ca) for further visualisation and functional analysis.

### Isolation of bacteria

For aerobic cultivation four media were chosen to culture different bacteria:

- Marine Agar (MA, BD Difco),
- R2A-SS (R2A (Merck) prepared with Red Sea Salt™ (RSS) at 36 ppt salinity),
- Supplemented Trypticase Soy Agar (TSA^+^ Merck supplemented with ferrous ammonium sulfate solution 5% ((NH_4_)_2_Fe(SO_4_)_2_, ThermoFisher), magnesium sulfate (MgSO_4_, Sigma) and sodium DL-lactate 60% (CH_3_CH(OH)COONa, Sigma), and,
- Supplemented Marine Agar (MA^+^, Marine Agar supplemented with (NH_4_)_2_Fe(SO_4_)_2_ solution 5% and sodium DL-lactate 60%)

The R2A-SS was autoclaved (121°C for 16 min) separately in 2× concentration of 15.2 g/L of R2A (pH 7.2 ± 0.2) and 36 g/L of RSS (pH 8.3 ± 0.2) to avoid salt precipitation, then mixed together prior to pouring into petri dishes. The MA (pH 7.6 ± 0.2) was prepared by sterilising 37.4 g/L of Marine Broth (BD, Difco) supplemented with 15 g/L of bacterial agar (BD, Difco). The TSA^+^ was prepared by adding 30 g/L of trypticase soy broth (BD, Difco), 0.98 g/L of MgSO_4_, 4 mL/L of sodium DL-lactate 60% and 15 g/L of agar (pH adjusted to 7.5 ± 0.05) and autoclaving. A 5% solution of (NH_4_)_2_Fe(SO_4_)_2_ was filter sterilised with a 0.22 μm pore size membrane and 20 mL added to 1 L of the cooled TSA+magnesium+lactate medium prior to pouring into petri dishes. The MA^+^ (with lactate and ferrous ammonium sulfate) was prepared by mixing MA and sodium DL-lactate (pH adjusted to 7.5 ± 0.05), autoclaving, adding the (NH_4_)_2_Fe(SO_4_)_2_ solution to the cooled MA, then pouring into petri dishes. All components were dissolved in distilled water.

For anaerobic cultivation, THA^+^ (Thioglycollate Agar, Merck (pH 7.1 ± 0.2) was prepared with RSS in the same fashion as R2A) was used instead of R2A-SS in the aerobic isolation, as it contains components designed to reduce the redox potential of the medium, favouring anaerobes. Thus, the four media used for anaerobic cultivation were MA, TSA^+^, MA^+^ and THA^+^.

Serial dilutions of samples from the inner and outer layers of one orange tubercle under which pitting had occurred were prepared in sterile saline (NaCl 0.9%). The mixtures were vortexed thoroughly for 1 min between dilutions. Volumes of 50 µL of four dilutions (undiluted and dilutions of 10^−1^, 10^−2^, 10^−3^) were spread plate inoculated onto prepared media in triplicate. Anaerobic conditions were achieved by an anaerobic jar (BD) containing anaerobic sachets (AnaeroGen, Oxoid) and indicators (Resazurin, Oxoid). All plates and anaerobic jars were incubated at 25°C for 16 days (aerobes) and for 25 days (anaerobes). All different colony morphologies from the growth on the plates were counted and subcultured onto the corresponding media to get pure isolates. The pure cultures were grown and stored in 20% glycerol (Merck) at –80°C as stocks.

### Identification of isolates

Isolated colonies of pure cultures were grown in liquid media with shaking at 150 rpm and on agar plates at 25°C for 48 h, which were then used for DNA extraction. Colony DNA (cDNA) was obtained by heat treatment of bacterial colonies in sterile MilliQ water at 95°C for 15 min. The mixture was rapidly cooled to ambient temperature and centrifuged at 10,000 *g* for 1 min to remove cell debris in the pellet. For those isolates where no cDNA was obtained, pure culture broths were used to extract gDNA by the MoBio UltraClean^TM^ Microbial DNA Isolation Kit (Qiagen), following the manufacturer’s instructions. The size and quality of extracted DNA was checked by electrophoresis in 1% agarose gel (Bioline) including GelRed™ (Biotium). DNA was quantified using Nanodrop. The 16S rRNA genes were amplified using primers 27F (5’-AGA GTT TGA TCM TGG CTC AG-3’) and 1492R (5’-CGG YTA CCT TGT TAC GAC TT-3’) (47). Each reaction tube contained 100–400 ng of template DNA (determined using Nanodrop), 12.5 µL of MangoMix (Bioline), 1 µL of each primer (10 µM), and sterile MilliQ water up to 25 µL. The run cycle of MyCycler™ (Biorad) for PCR was: an initial denaturation at 95°C for 5 min; 30 cycles of denaturation (95°C for 45 sec), annealing (55°C for 45 sec), and extension (72°C for 45 sec); and a final extension at 72°C for 5 min. The PCR products were sent to Macrogen Inc. (Korea) in a 96 well-plate at ambient temperature for DNA purification and Sanger sequencing. Received sequence chromatograms were viewed in the software package BioEdit and manually checked for quality and length. Trimmed high-quality sequences were compared to those in GenBank at the National Center for Biotechnology Information (NCBI) by the nucleotide Basic Local Alignment Search Tool (BLASTn). Sequences with ≥97% identity to the isolate sequences were deemed putative identifications of the isolates. 16S rRNA gene sequences of type strains of the isolate’s closest relatives were obtained from the List of Prokaryotic names with Standing in Nomenclature (LPSN). Phylogenetic trees were constructed in MEGA7.0.14 (Molecular Evolutionary Genetics Analysis software, www.megasoftware.net) using Clustal W for sequence alignment and the Neighbour Joining clustering method with 2000 bootstrap resamplings.

### Accession numbers

The microbial sequencing data have been deposited on NCBI. The bioproject accession number PRJNA524055 (https://www.ncbi.nlm.nih.gov/bioproject/PRJNA524055) has been issued for the metabarcoding dataset (9 biosample accessions from SAMN10995695 to SAMN10995703). For isolates’ sequences, the accession numbers (https://submit.ncbi.nlm.nih.gov/subs/genbank/SUB5221871) have been continuously from MK568337 to MK568455 which are included in Fig. 7, next to corresponding isolate codes.

## ACKNOWLEDGMENTS

Hoang C. Phan acknowledges the support of Victorian Government (Australia) via a joint scholarship between the Victorian International Research Scholarship and Swinburne University of Technology. The authors would like to thank Sean Blackwood (Queenscliff Harbour) for assisting with field testing and providing samples of sheet piling steels used in this work.

